# No detectable DNA methylation in the kelps (Laminariales): a conversion-free test of a predicted loss

**DOI:** 10.64898/2026.11.17.767002

**Authors:** Robin S. Sleith, Scott Lindell, Michael W. Lomas, Gary Molano, Nichole N. Price, Charlotte T. C. Quigley, Peter D. Countway

## Abstract

Kelps (Laminariales) dominate temperate and polar rocky coasts and support a fast-growing aquaculture sector. Breeding and restoration programs need to know whether a line’s performance follows its genotype or its culture history. DNA methylation has been reported in kelps, with differences by life stage, thermal history and population of origin now underpinning a literature on conditioning and priming in seaweed breeding. Recent comparative genomics work predicts otherwise, because brown algae lack the canonical DNA methyltransferases and should therefore carry little or no DNA methylation. We tested the prediction with native-DNA nanopore sequencing, which requires neither bisulfite conversion nor methylation-dependent digestion, in seven kelp datasets spanning three genera, four species and four *S. latissima* life stages. Nuclear 5-methylcytosine stayed at the measurement background in every sequence context (CG 0.021-0.034% of sites, against 0.023% on PCR copies that carry no methyl group by construction), while the same pipeline returned 58% of CG sites methylated in a diatom and recovered the transposon-targeted distribution a functioning system predicts. In native reads, thirteen of the fourteen best-covered chloroplast positions that bisulfite sequencing placed above 2% carry at most one methylated call, and those published levels track each library’s conversion efficiency rather than its tissue. Gene content and direct measurement agree, so the heritable effects reported in cultivated kelp require a carrier other than DNA methylation: chromatin modifications or non-coding RNA.

---

Kelps (Laminariales) form the dominant canopy of temperate and polar rocky coasts, where they support productive and diverse communities, and they are also the basis of a fast-growing aquaculture sector and of restoration efforts on coasts where forests have been lost or degraded. Both uses place the same demand on kelp biology: an understanding of which traits are heritable, and how reliably they are transmitted from one generation to the next.

Breeding programs for cultivated and restoration kelp now select among gametophyte lines and crosses on performance measured in common gardens and on farms, and increasingly use genomic tools to do so (Lindell, 2026; DeWeese et al., 2025). Seed stock passes through clonal gametophyte cultures that may be held for years before outplanting, which makes the separation of genotype from environmental history unusually difficult: a line’s performance may reflect its alleles, its culture history, or an interaction between the two. A molecular mechanism that carries environmental experience across cell divisions, and across the alternation of generations, would therefore matter both for how cultivars are selected and for how common-garden results are read.

Epigenetic regulation in brown algae has been studied in both evolutionary and applied work. Histone modifications have been mapped through the life cycle of the model brown alga *Ectocarpus* (Bourdareau et al., 2021), chromatin state has been linked to sexual differentiation in its UV sex determination system (Gueno et al., 2022), and chromatin accessibility has been linked to generation-biased gene expression in *Saccharina japonica* (Yang et al., 2025). DNA methylation has received the most attention, being the best-characterized heritable chromatin mark in other eukaryotes, and the picture it presents is contradictory. The first brown algal genome found no detectable cytosine methylation by two conversion-free assays, and attributed the silencing of its transposons to small RNAs instead (Cock et al., 2010). Bisulfite sequencing of *Saccharina japonica* subsequently placed 5-methylcytosine at approximately 1.4% of cytosines, with differences among male gametophytes, female gametophytes and young sporophytes (Fan et al., 2020), and reported methylation of the chloroplast genome in the same libraries (Teng et al., 2021); later work reported a response to heat stress (Liu et al., 2023) and, using methylation-dependent digestion, differences by population of origin and cultivation history in *S. latissima* (Scheschonk et al., 2023, 2024). These values now underpin an applied literature on hatchery conditioning, thermal priming and the prospect of epigenetic selection in seaweed breeding.

Comparative genomics predicts otherwise. Cytosine methylation is placed on DNA by a small and well-conserved set of enzymes (de Mendoza et al., 2020): DNMT1 maintains the mark by copying it to the new strand at each round of replication, and DNMT3-type enzymes establish it anew. Many algae and some fungi instead rely on DNMT5, an ATP-dependent enzyme that maintains CG methylation in the fungus *Cryptococcus neoformans* (Catania et al., 2020; Dumesic et al., 2020). In the diatom *Phaeodactylum tricornutum*, knocking out DNMT5a depletes CG methylation across the genome and de-represses transposons (Hoguin et al., 2023). A fourth member of the family, DNMT2, is widely conserved but is not a DNA methyltransferase: wherever its activity has been characterized it methylates a cytosine in transfer RNA (Jeltsch et al., 2017). A genome that encodes none of the DNA-methylating enzymes has no known means of methylating its own DNA, so the presence or absence of these genes is informative in itself. DNMT1 is present in the earliest-branching brown alga and in non-phaeophycean relatives but nowhere else in the Phaeophyceae, from which it was anticipated that most brown algae would lack DNA methylation or carry very little (Denoeud et al., 2024), and a census of chromatin proteins across the clade found no DNMT1, DNMT3 or DNMT5 in any species examined (Vigneau et al., 2026). Neither study measured DNA methylation in a kelp, so the conflict between the reported methylomes and the absent machinery has stood unresolved.

Resolving it calls for a method whose errors are independent of those assays and, more importantly, can be measured within the same sample. Bisulfite conversion is monitored with a spiked-in unmethylated control, but that template is foreign to the sample and unstructured, so it reports the average conversion of the reaction rather than conversion at any particular position in the genome of interest (Warnecke et al., 2002; Genereux et al., 2008); methylation-dependent digestion recovers only modified fragments by construction (Wang et al., 2015), and so admits no unmethylated control at all. Native-DNA nanopore sequencing (Oxford Nanopore) meets that condition. Modifications are inferred from the ionic current of unmodified template, so no conversion or digestion step intervenes; its calls carry their own error, which we do not claim to be smaller, but that error can be quantified from unmethylated DNA carried through the same chemistry, as was done to settle an equivalent question about human mitochondrial DNA (Bicci et al., 2021). In every sample reported here, that control is the organism’s own plastid genome. Here we report direct, conversion-free measurements of 5mC and 6mA in seven kelp datasets spanning three genera and four species of Laminariales, including four life stages of *S. latissima*, each against its own chloroplast as an internal control carried through the same chemistry and against a methylated diatom as a positive control, and we re-examine, in native reads of the species in which they were reported, four chloroplast cytosines that bisulfite sequencing placed at around 7% methylated.

Seven kelp datasets and one diatom positive control were processed through a single pipeline. Gametophyte cultures of *Saccharina latissima* of both sexes, a meiospore pool and sporophyte blade tissue, cultures of *S. japonica* (CCMP10001) and *Macrocystis pyrifera* (CCMP10000) from the National Center for Marine Algae and Microbiota, and field material of *Alaria esculenta* were sequenced on R10.4.1 flow cells using the ligation kit SQK-LSK114 and basecalled with Dorado 1.3.4 (sup@v5.2.0 with 5mC_5hmC@v2 and 6mA@v1). The same molecules were re-basecalled with HAC v6.0.0 (5mC_5hmC@v1 and 6mA@v1, Dorado 2.1.2) and filtered identically, so that individual calls can be tested for dependence on the model.

Nuclear assemblies were those of DeWeese et al. (2025) for *S. latissima*, Diesel et al. (2023) for *M. pyrifera*, Bringloe et al. (2022) for *A. esculenta*, and GCA_059791855.1 for *S. japonica*. Each was screened for bacterial sequence and for organelle-like contigs, then combined with the chloroplast and mitochondrial genomes of its own species so that organelle reads map competitively to the organelles, and species identity was confirmed for every sample from its own organelle reads. Alignment was with minimap2 2.28 (Li, 2018) (-ax map-ont -y, carrying modification tags), retaining primary alignments at MAPQ >= 20 with at least 75% of the read aligned, and all levels were then computed on reads at least 95% identical to the reference (the identity filter); values computed without it are given where stated. A stricter 98% cut removes 11 to 31% of nuclear reads against 3 to 15% of chloroplast reads; splitting reads into identity bands shows that those between 95 and 98% carry more modification calls in both compartments, so the reads it discards hold no hidden nuclear methylation. Calls were made with modkit 0.6.4 pileup at a fixed threshold of p >= 0.8 for both C and A, separately for CG, CHG and CHH, with call-confidence distributions from modkit sample-probs. Per-read and per-segment CG levels were scored directly from the MM/ML modification tags under the same rule, and 2-kb read segments were classified by DIAMOND blastx against RepeatMasker transposable-element proteins or the species’ gene catalog at e <= 1e-5.

Each sample’s own chloroplast, sequenced on the same flow cell and called by the same model, is the within-sample reference for its nuclear values. Plastid DNA is reported as unmethylated in most land plants but is hypermethylated in *Chlamydomonas* maternal gametes by a chloroplast-resident methyltransferase (Nishiyama et al., 2002), so we assume nothing about its state: any methylation it carried would raise the reference and make the nuclear comparison more conservative. A PCR product provides a second reference that carries no methylation by construction: a full-length 16S amplicon run (27F/1492R) of the same *Alaria* holdfast material, sequenced on the same flow cell as the *Alaria* metagenome, from which the 150 reads matching the *A. esculenta* mitochondrial small-subunit rRNA gene (NC_058765, at least 97% identity and at least 1% above any other kelp organelle genome) were called on that gene. The diatom *Thalassiosira* serves as the methylated positive control, from a pooled coastal plankton sample collected in Maine during a June 2026 *Pseudo-nitzschia* event and mapped to *Thalassiosira delicatula* AJA262-20 (GCA_036939835.2, isolated at Boothbay Harbor) with its plastome. For the chloroplast positions, organelle reads were assigned to species by competitive alignment to six kelp organelle genomes and *S. japonica* plastid reads were called with modkit extract, alongside reprocessing of the Fan et al. (2020) bisulfite libraries (SRR7528100-108) with Bismark (Krueger & Andrews, 2011) against the same plastid plus lambda. Sample provenance, read counts, coverage and the full per-context values are given in Table S2.

Nuclear 5mC was indistinguishable from background in every kelp sample, genus and sequence context (Figure 1a). The share of nuclear CG sites called modified ranged from 0.021% in *S. japonica* to 0.034% in the male *S. latissima* gametophyte, in every case below the same sample’s own chloroplast (0.032 to 0.047%); CHG and CHH behaved the same way, at 0.059 to 0.089% and 0.067 to 0.095% nuclear against higher plastid values. Because the chloroplast was sequenced on the same flow cell and called by the same model, this is a measured floor rather than an assumed one, and no kelp nuclear value rises above it. A second, independent floor comes from DNA that cannot be methylated: PCR copies of *Alaria*’s own mitochondrial rRNA gene, sequenced alongside the metagenome, on which every call is a false positive. *Alaria* nuclear CG, CHG and CHH (0.025, 0.066 and 0.079%) sit within that amplicon’s level or its 95% upper limit in every context (amplicon 0.023, 0.032 and 0.087% on 150 reads; the CG and CHG rates rest on two and three methylated calls, upper limits 0.083 and 0.095%; Table S3). The distribution of call probabilities is consistent with the same interpretation: kelp nuclear 5mC calls carry the low-confidence profile of their own chloroplast calls, with no accumulation of calls near certainty (Figure 1b). These residual levels are what independent benchmarking of these callers predicts, since in genomes with little non-CG methylation the error is dominated by false-positive rather than false-negative calls and falls further at thresholds stricter than the p >= 0.8 applied here (Kulkarni et al., 2026). Within *S. latissima* the measurement spans the life cycle. Haploid female and male gametophyte cultures, a pool of meiospores released from fertile sporophytes, and diploid sporophyte blade tissue all give the same result: nuclear CG between 0.024 and 0.034% of sites, in every case below that sample’s own chloroplast (0.039 to 0.056%), with CHG and CHH behaving likewise. The blade matches the gametophytes once the identity filter is applied; without it the blade reads higher in nucleus and chloroplast alike, because its reads are the least accurate of the four (mean identity 96.6%). The meiospore pool, the deepest of these datasets, lies in the same range. Life stage is the comparison in which the published methylomes reported their largest differences, and across these four stages we find none. Because spores and blade tissue mix cell types, methylation confined to a subset of cells would be diluted in a genome-wide average while still appearing as a tail of heavily methylated reads, as it does in the diatom. In 200,000 random nuclear reads per stage, scored without the identity filter, there is no such tail: of 314,846 reads with normal call density, one reaches 10% CG methylation, in the blade. A methylated subpopulation above roughly 0.003 to 0.02% of nuclear reads, depending on stage, would have been detected.

**FIGURE 1.**
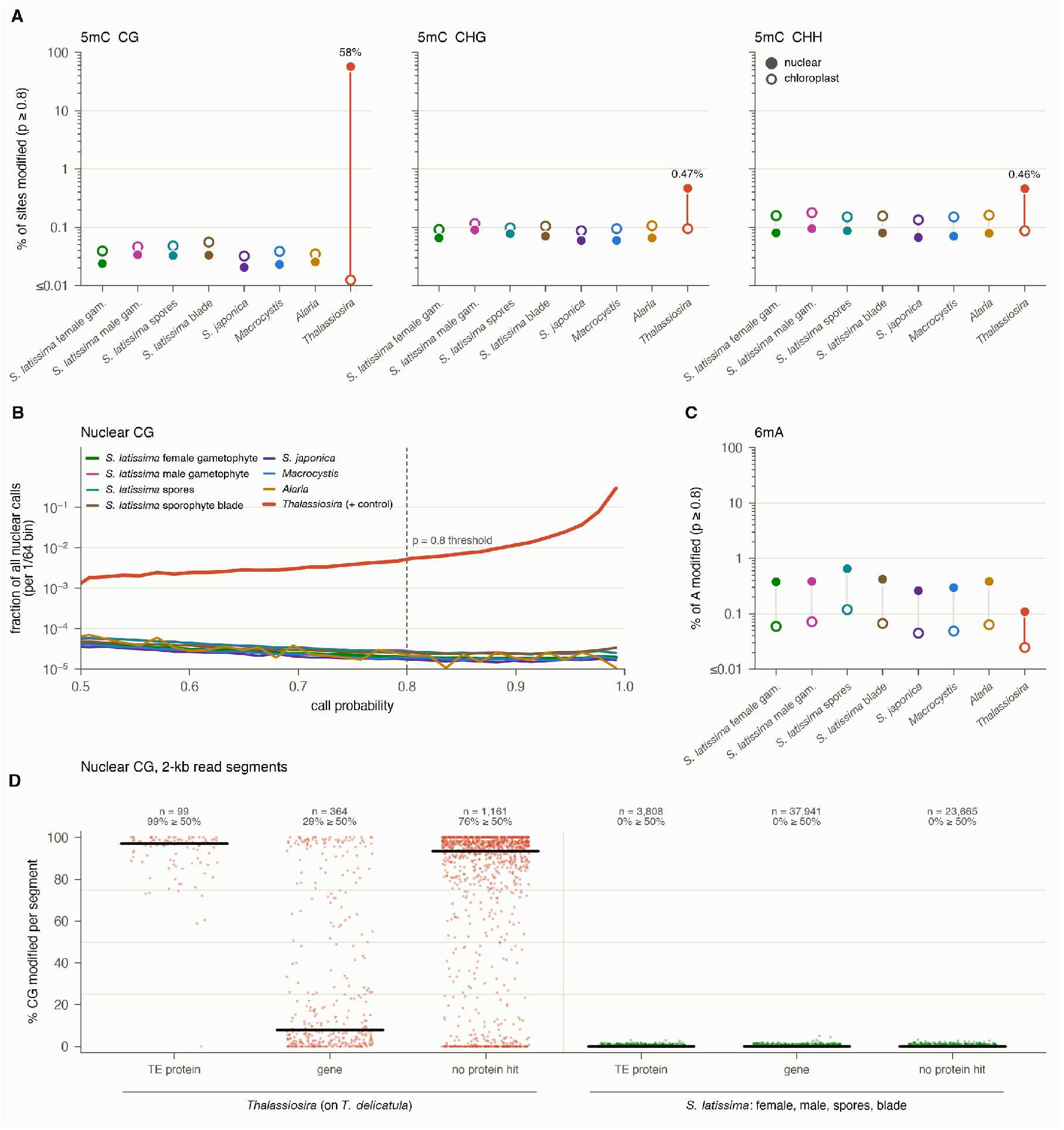
All samples were basecalled with the same model and filtered identically, on reads at least 95% identical to the reference. (a) Percentage of sites called 5mC in CG, CHG and CHH; filled symbols nuclear, open symbols each sample’s own chloroplast, sequenced on the same flow cell and called by the same model. *S. latissima* appears at four life stages. (b) Confidence of nuclear CG calls whose top call is 5mC. (c) Percentage of adenines called 6mA, on the same layout. (d) CG methylation of 2-kb nuclear read segments classified by protein homology as transposable-element, genic or unmatched, with the diatom mapped to *Thalassiosira delicatula* and *S. latissima* pooled across the four life stages; black bars are medians.

The same pipeline detects methylation where it exists. Reads assigned to the diatom *Thalassiosira* return 57.5% of nuclear CG sites modified, three orders of magnitude above any kelp value, with its own chloroplast at 0.012%, and its calls pile up above p = 0.95 where every kelp distribution stays flat (Figure 1a,b). The diatom signal is also distributed as a functioning system predicts rather than diffusely: 98 of 99 2-kb read segments carrying a transposable-element protein match are methylated at half or more of their CG positions, against 29% of 364 segments matching genes, while in *S. latissima* none of 3,808 transposable-element segments reaches that level, across all four life stages (Figure 1d), consistent with the unmethylated transposons of the *Ectocarpus* genome (Cock et al., 2010). The signal is carried by the reads themselves rather than by reference fit: 61% of diatom nuclear reads carrying at least 50 CG calls are half or more methylated (608 of 992), and no chloroplast read is. Reads whose best match among the candidate genomes is the independent *T. rotula* FE80 assembly are 71.1% methylated (256 reads), against the 69.2% measured by native nanopore sequencing in a culture of that same strain (Di Costanzo et al., 2025). Within the kelp libraries themselves the GANTC adenines of co-sequenced bacteria are called modified in a majority of reads, confirming that a genuinely modified base in these samples is recovered.

The gene content of the species accounts for the measurement (Table 1). The published census covers *S. latissima* but not *S. japonica*, the species in which the methylomes were reported, and scores neither DNMT2 nor DNMT5 (Vigneau et al., 2026), so we searched the genomes of all four species directly. Each encodes a single C5 cytosine methyltransferase, carrying the methyltransferase domain alone and falling with DNMT2, the transfer-RNA enzyme; no DNMT1, DNMT3, DRM, chromomethylase or DNMT5 was recovered, nor any MT-A70 adenine methyltransferase, while control queries in the same searches found their expected targets. Two hits cleared the cutoff without being methyltransferases: one matches only DNMT5’s helicase segment, on a 559-bp contig at 61% GC, and the other only DNMT1’s CXXC zinc finger, within a 1,936-residue protein lacking any methyltransferase domain. This matches the brown algal pattern reported from genome surveys of the clade (Denoeud et al., 2024). A recent review proposes that DNMT2 compensates for the loss of DNMT1 and maintains the reported methylation in brown algae (Khatei et al., 2026); DNMT2-family enzymes act on transfer RNA in the organisms where they have been characterized, and we find no DNA methylation for such an activity to maintain.

The chloroplast methylation reported for *S. japonica* can be examined directly, because the same positions can be read in native DNA of the same species, and the plastid sequences are effectively identical, so any difference is not one of strain. The deposited tables place 47 chloroplast cytosines at 2% or above in gametophytes; the fourteen that rest on at least a thousand reads all lie in predicted stable stems, and in native reads thirteen of them carry at most one methylated call, three in 4,578 pooled, with every upper limit below the published level (Figure 2a). Among them are the four cytosines of the rbcL-rbcS spacer hairpin, placed at 6 to 7% by bisulfite sequencing, which carry one of 1,557 native calls (Figure 2c; Figure S1). The single exception, at 117,907, is called by one basecalling model and not another, sits in a homopolymer run that the caller over-calls, and has a stem partner of identical sequence that reads none of 417. The published levels follow conversion rather than tissue. Re-processed library by library, the plastid-wide level equals the unconverted fraction of the lambda spike-in, and the hairpin reads 6.5 to 9.5% in the six gametophyte libraries, whose lambda conversion was the poorest, against 0.03 to 0.85% in the three better-converted sporophyte libraries (Figure 2b). Methylation would not track conversion efficiency; incomplete conversion of stable stems would. We take this as illustrating a general hazard rather than a fault of those studies: where true methylation is below one per cent, the residual of bisulfite conversion is of the same size as the signal and is concentrated in exactly such structured regions, while methylation-dependent digestion cuts only at modified cytosines, so that every recovered tag is methylated by construction. These are the conditions under which a low apparent level is difficult to distinguish from none at all.

**FIGURE 2.**
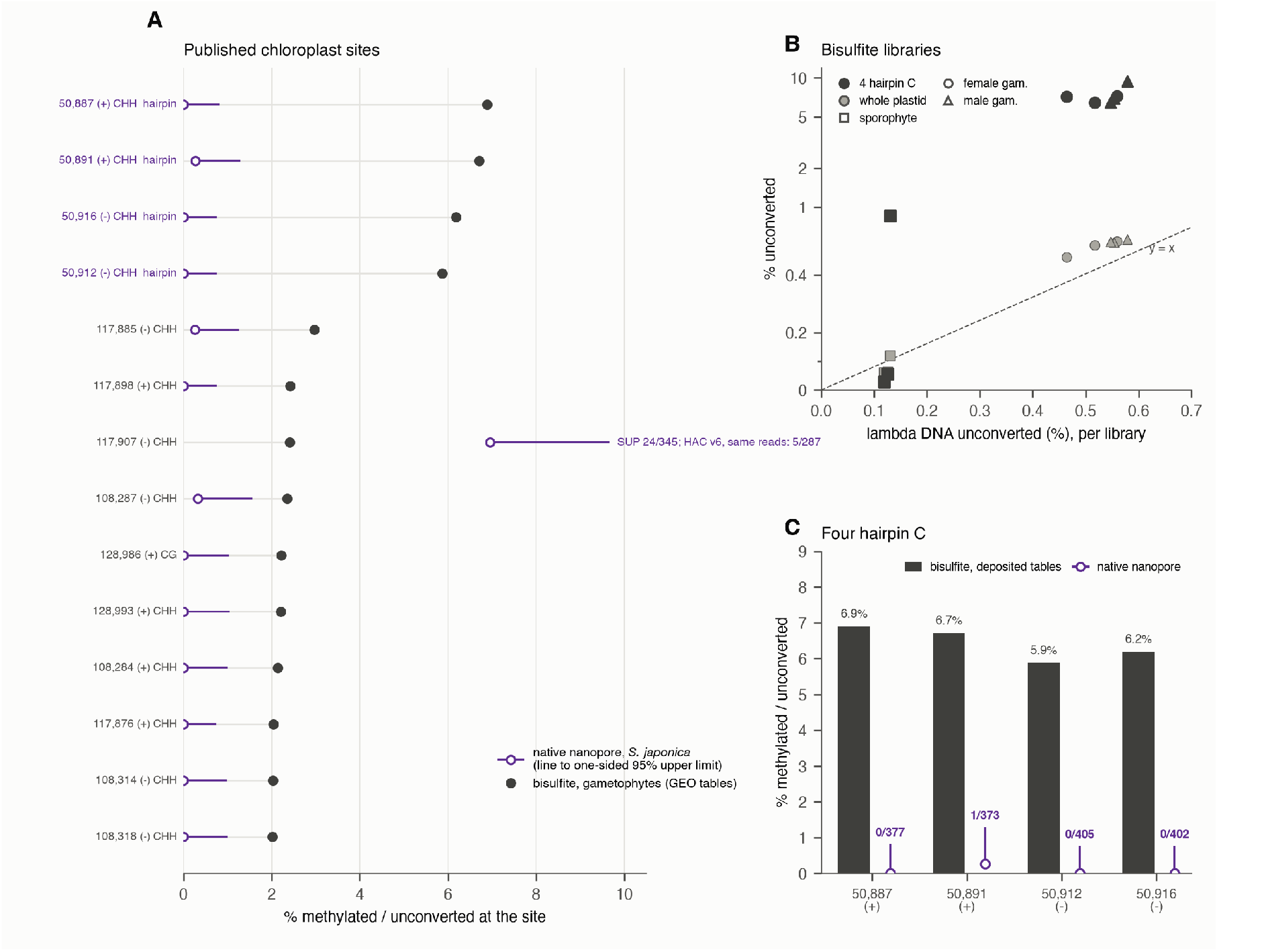
Published chloroplast sites read in native DNA. (a) Every chloroplast cytosine that the deposited bisulfite tables place at 2% or above in gametophytes on at least a thousand reads (14 sites), ranked, against the native level at the same position with its one-sided 95% upper limit. A further 33 sites reach 2% on three to nine unconverted reads, and the six with at least a hundred native calls give one methylated call in 1,399. (b) The Fan et al. (2020) libraries re-processed: the whole-plastid level and the four hairpin cytosines against lambda non-conversion, library by library. (c) The four hairpin cytosines, deposited bisulfite tables against native counts.

Nuclear 6mA is the one measurement that exceeds the plastid floor (Figure 1c), and two artefacts account for the difference. Most of the nuclear positions called 6mA in a substantial fraction of reads lie on a single scaffold of the *S. latissima* assembly (DeWeese et al., 2025; scaffold_208, 175–914 kb), where two complete plasmids of *Pseudosulfitobacter pseudonitzschiae* (99.9% identical to strain SMR1, CP022416–CP022417) are joined to kelp sequence, carrying that bacterium’s own GANTC methylation; after masking the bacterial sequence, few remain and none is strongly supported. The positions that survive the mask are caller background. The rate at which adenines are called modified depends on the base composition of the read, rising about tenfold from the most GC-poor to the most GC-rich reads, and it does so to the same degree in nuclear and in plastid DNA. Nuclear reads are much the GC-richer of the two, and matching the compartments on GC accounts for the difference between them. In the subset of reads aligning at 98% identity or better, stricter than the 95% used elsewhere and chosen so that GC bins can be populated in both compartments, the nuclear rate predicted from the plastid at the same GC is 0.26% against 0.25% observed in the female gametophyte and 0.25% against 0.23% in the male; across four samples the observed rate falls between 0.6 and 1.2 times the prediction, where the unmatched comparison differs four-to six-fold. Rates in that subset sit below the whole-pileup values and are not directly comparable with them. We therefore treat kelp 6mA as background; no adenine methyltransferase of the MT-A70 family is present in any of the four genomes.

Taken together with the gene content of the clade, these measurements bring three bodies of work into agreement. The first brown algal genome reported no detectable cytosine methylation and proposed small RNAs as the silencing mechanism (Cock et al., 2010); comparative genomics later found the canonical methyltransferases absent across the Phaeophyceae and predicted that methylation would be lost with them (Denoeud et al., 2024; Vigneau et al., 2026); and we find none, at base resolution, in four species of three kelp genera. The published kelp methylomes stand apart from this consensus, and our data indicate that the difference lies with the assays. Two consequences follow. The first is methodological and applies to any low-methylation genome: a measurement near one per cent needs unmethylated sequence from within the sample itself, not a spiked-in template, because a spike reports the average performance of the reaction while at any individual position a conversion floor and a biological signal look alike. Native reads supply such a reference in every sample; here it is the plastid, whose own level is measured rather than assumed, and any methylation it carried would only make the comparison more conservative. The second is for the applied literature, and is specific rather than deflationary: gametophyte thermal priming is reported to alter the transcriptomic heat-stress response of sporophyte progeny (Nilsen et al., 2026), and those effects are not in question here, only the carrier. The candidates that remain are the ones both the earliest and the most recent work point to, chromatin modifications and small or other non-coding RNAs, and we suggest that effort move there.

## Abbreviations

5mC: 5-methylcytosine
5hmC: 5-hydroxymethylcytosine
6mA: N6-methyladenine.

---

**FIGURE S1.**
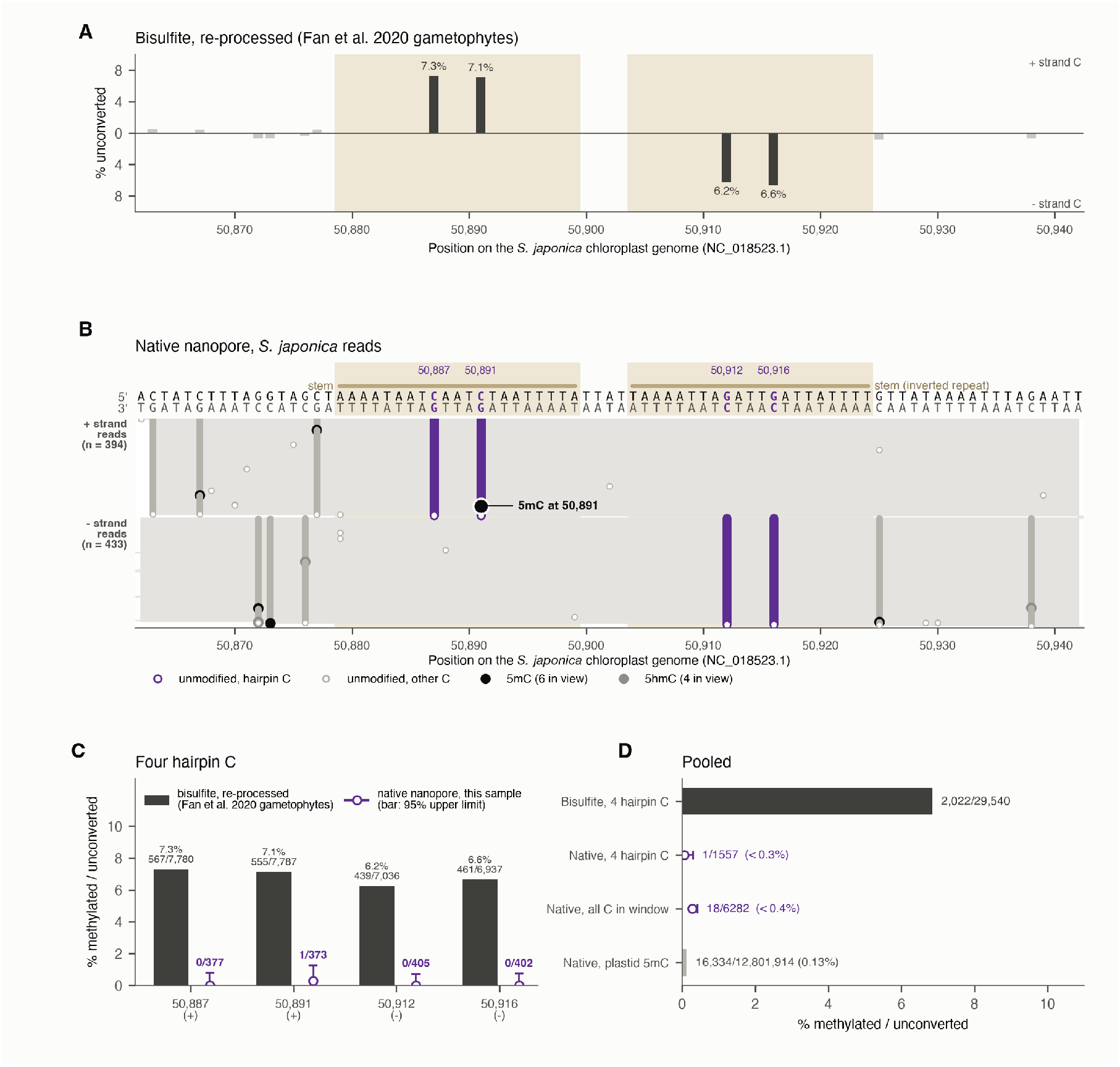
The hairpin in detail. (a) Fan et al. (2020) bisulfite data for gametophytes, re-processed from the raw reads, at every cytosine in the window; the four hairpin cytosines are the stem cytosines. (b) Every native cytosine call at p >= 0.8 on S. japonica reads at least 95% identical to the reference crossing the hairpin, one row per read. (c) The four hairpin cytosines, re-processed bisulfite against native. (d) Pooled levels for the hairpin, the surrounding window and the whole plastid.

## ACKNOWLEDGMENTS

We thank Brittney Honish for assistance in the laboratory, Owen Smith in the laboratory and the field, and Sarah Douglas in the field; Sydney Greenlee for processing the diatom sample; and Tyler Spillane and Bryant Lewis for providing the Maine Department of Marine Resources plankton sample. This work was supported by the World Wildlife Fund (grant number [TO ADD]).

## Notes

### Competing Interest Statement

The authors have declared no competing interest.

